# Separation of phosphatase and kinase activity within the Bub complex is required for proper mitosis

**DOI:** 10.1101/2022.04.11.487869

**Authors:** Lei Wang, Thomas Kruse, Blanca López-Méndez, Yuqing Zhang, Chunlin Song, Lei Zhu, Bing Li, Jing Fang, Zhimin Lu, Jakob Nilsson, Gang Zhang

**Affiliations:** The Cancer Institute, The Affiliated Hospital of Qingdao University, Qingdao University, Qingdao, China; Novo Nordisk Foundation Center for Protein Research, Faculty of Health and Medical Sciences, University of Copenhagen, Copenhagen, Denmark; The Department of Genetics and Cell Biology, Basic Medical College, Qingdao University, Qingdao, China; Institute of Translational Medicine, Zhejiang University School of Medicine, Hangzhou, China

## Abstract

The Bub1 and BubR1 kinetochore proteins support proper chromosome segregation and mitotic checkpoint activity. Bub1 and BubR1 are paralogues with Bub1 being a kinase while BubR1 localizes the PP2A-B56 protein phosphatase to kinetochores in humans. Whether this separation of kinase and phosphatase activity is important is unclear as some organisms integrate both activities into one Bub protein. Here we engineer human Bub1 and BubR1 proteins integrating kinase and phosphatase activities into one protein and show that these do not support normal mitotic progression. A Bub1-PP2A-B56 complex can supports chromosome alignment but results in impairment of the checkpoint due to dephosphorylation of the Mad1 binding site in Bub1. Furthermore, a chimeric BubR1 protein containing the Bub1 kinase domain induces delocalized H2ApT120 phosphorylation resulting in reduction of centromeric hSgo2 and chromosome segregation errors. Collectively, these results argue that the separation of kinase and phosphatase activities within the Bub complex is required for balancing its functions in the checkpoint and chromosome alignment.

## INTRODUCTION

The accurate segregation of the genetic material during cell division requires that kinetochores establish proper connections to the microtubules of the mitotic spindle. A complex surveillance mechanism, the spindle assembly checkpoint (SAC), monitors kinetochore-microtubule interactions and delays mitotic exit until all kinetochores have bound to microtubules^1-3^. The SAC is composed of a set of conserved checkpoint proteins that associates with outer kinetochore proteins during prometaphase to generate a “wait anaphase” signal. Generation of the SAC signal depends on the assembly of a Mad1/2-Bub1 complex, which is mediated by Mps1 phosphorylation of Bub1 Thr 461 in humans^4-9^. In turn, the Mad1/2-Bub1 complex stimulates the generation of the mitotic checkpoint complex (MCC) composed of the BubR1 and Mad2 checkpoint proteins bound to Cdc20^5, 6, 10-12^. The MCC constitute the biochemical identity of the “wait anaphase” signal.

In addition to generating a checkpoint signal in response to unattached kinetochores several checkpoint proteins facilitate the establishment of proper kinetochore-microtubule interactions. In particular, the Bub1 and BubR1 checkpoint proteins have been shown to be important for alignment of chromosomes independent of their checkpoint function^13-16^. Bub1 and BubR1 are paralogues that arose from gene duplications of an ancestral “MadBub” gene^17-19^. Bub1 and BubR1 share a common GLEBS interaction motif for Bub3 and directly bind each other through a region C-terminal to the Bub3 binding region (Fig. 1A)^20-22^. In addition, numerous short linear motifs (SLiMs) are present throughout the Bub1 and BubR1 proteins that mediate interactions with specific mitotic regulators (Fig. 1A)^13^. At the C-terminus both proteins contain a kinase domain that in the case of BubR1 is a pseudokinase domain^23^ while Bub1 is an active kinase that phosphorylates T120 on histone H2A to facilitate hSgo1 and hSgo2 recruitment to centromeres^24, 25^.

**Figure 1.**
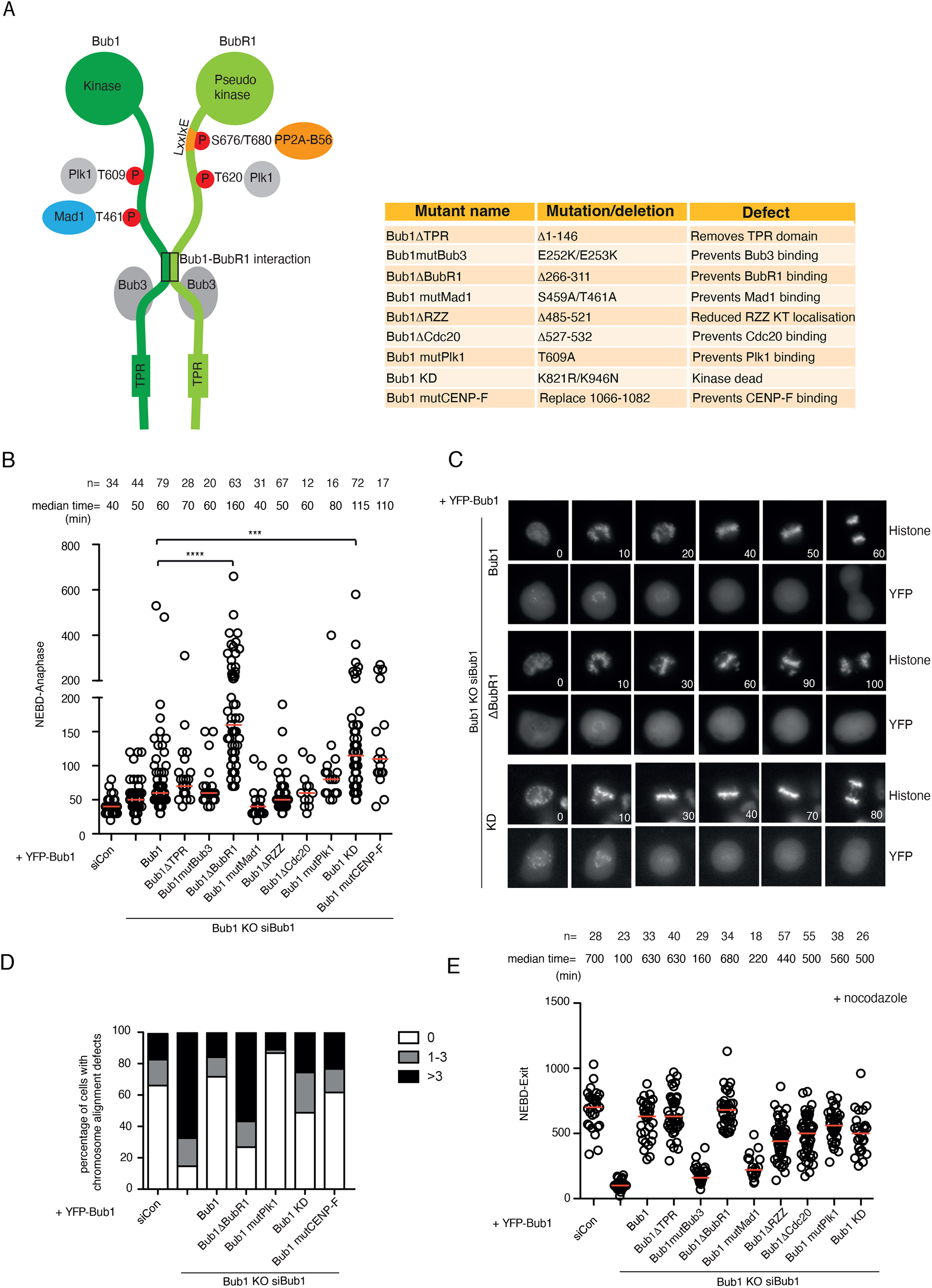
Systematic analysis of Bub1 interaction modules. **A**) Schematic of the Bub1-BubR1 complex and phosphorylation sites and interactions and table of the mutants of Bub1 analysed. **B**) The time from NEBD to anaphase in cells complemented with the indicated YFP-tagged Bub1 constructs. Each circle represents the timing of a single cell and the red line indicates the median time. Mann-Whitney u-test was applied. ns means not significant. **** means P<0.0001, *** means P<0.001. The number of cells analysed for each condition is indicated on top (n=X). Representative experiment of at least two independent experiments of each mutant. **C**) Representative still images of mitosis in cells complemented with Bub1, Bub1ΔBubR1 and Bub1 KD. The top panel is CFP-H3 and the bottom panel is YFP-Bub1. **D**) Quantification of the chromosome alignment defects in cells complemented with the indicated Bub1 constructs. Cells were stained with DAPI and antibodies for tubulin, YFP and CENP-C and number of unaligned chromosomes counted. 100 cells were analysed per condition and the experiment performed twice and results from one of the experiments is shown. **E**) The time from NEBD to mitotic exit of a single cell in nocodazole treated cells complemented with the indicated Bub1 constructs. Each circle represents the time from NEBD to mitotic exit and the red line indicates the median time. The number of cells analyzed per condition is indicated above (n=X). Representative experiment of at least two independent experiments of each mutant.

BubR1 supports chromosome alignment by recruiting the PP2A-B56 Ser/Thr protein phosphatase through a well characterized LxxIxE motif to counterbalance Aurora B and Plk1 kinase activity at kinetochores^15, 22, 26-29^. Several functions of Bub1 in chromosome alignment have also been uncovered including kinetochore recruitment of Plk1 and CENP-F, binding to BubR1 as well as Bub1 kinase activity^13, 30-33^. However, the relative contribution of the many Bub1 interactions to chromosome alignment and SAC function is unclear. This is in part because investigation of Bub1 function has been challenging in human cells as very low levels of Bub1 is sufficient for its function^34-36^. We recently developed an approach that circumvents this problem by combining CRISPR/Cas9-mediated knockout of Bub1 and RNAi depletion of residual Bub1 allowing functional investigations into Bub1^34^. We here use this approach to perform a systematic analysis and side by side comparison of the different functional domains and SLiMs in Bub1 and their contribution to chromosome alignment and SAC function. This provided novel insights into Bub1 function and revealed a requirement for separation of kinase and phosphatase activities in the Bub1-BubR1 complex for accurate chromosome segregation in human cells.

## RESULTS AND DISCUSSION

### A systematic analysis of Bub1 functional modules

We recently used CRISPR/Cas9 to target Bub1 in HeLa cells hereby generating a cell line with low levels of endogenous Bub1^34^. Depletion of the residual Bub1 by RNAi results in almost complete removal of Bub1 with penetrant phenotypes in chromosome alignment and SAC signaling. By complementing Bub1 depleted cells with an RNAi resistant YFP-tagged Bub1 construct these phenotypes are suppressed providing an opportunity for analyzing the effects of Bub1 mutations in a clean null background. Numerous functional protein-protein interaction modules in Bub1 have been reported to regulate SAC signaling and chromosome alignment (Fig. 1A). However, these modules have not all been analyzed in a clean Bub1 null background making unambiguous comparisons difficult. Therefore, we decided to do a systematic side-by-side analysis of the modules of Bub1 to investigate their respective contribution to Bub1 function.

We combined cell synchronization with Bub1 RNAi depletion and complementation with YFP-tagged Bub1 constructs and followed mitotic progression by time-lapse microscopy. We recorded time from nuclear envelope breakdown (NEBD) to anaphase in single cells and monitored chromosome alignment using a fluorescent histone variant (Fig. 1B-C). From this analysis the disruption of three functional modules in Bub1 increased mitotic timing and induced chromosome alignment defects: the region binding to BubR1, the Bub1 kinase activity and the C-terminal helix reported to bind CENP-F. The important role of these functional modules was confirmed by immunofluorescence analysis of chromosome alignment in cells treated with a proteasome inhibitor (Fig. 1D). However, subsequent in-depth analysis of our Bub1 CENP-F mutant, which replaced the C-terminal helix in Bub1 with an alternative helix, revealed that it was defective in kinase activity, and we therefore did not investigate this mutant further (Supplemental Fig. 1A-B). We note that the GLEBS domain is essential for Bub1 function but due to the lack of a functional checkpoint in this mutant mitotic timing is not increased.

To analyze the effect on SAC signaling we used a similar experimental set-up and challenged cells with nocodazole to activate the checkpoint and recorded time from NEBD to mitotic exit (Fig. 1E). This revealed a strong requirement for the Bub3 interaction domain as well as the Mad1 interaction domain as expected. We observed only a minor contribution to SAC signaling from the ABBA motif that binds Cdc20^37^ suggesting that this interaction is not essential for the SAC in human cells in contrast to *C. elegans* and *in vitro* biochemical reconstituted assays^10, 11^.

In summary we have provided a careful analysis of the functional determinants of Bub1 revealing their relative contribution to chromosome alignment and SAC signaling. Bub1 interactions to Bub3 and Mad1 seemed to be the most important for SAC signaling whereas the strongest chromosome alignment phenotypes were observed when binding between Bub1 and BubR1 was uncoupled or Bub1 kinase activity disrupted.

### Chromosome alignment is supported by engineered Bub1 proteins binding PP2A-B56

Based on the above results we decided to investigate the role of the BubR1-Bub1 interaction in more detail. Since it is well established that BubR1 recruits PP2A-B56 to facilitate proper kinetochore-microtubule interactions we speculated that the lack of kinetochore localized PP2A-B56 in Bub1ΔBubR1 complemented cells was the cause of alignment defects. Consistent with this idea we observed increased kinetochore phosphorylation in Bub1ΔBubR1 complemented cells (Supplemental Fig. S1C). However, human Bub1 also contains a putative PP2A-B56 binding motif (aa 654-FSPIQE-659) and Bub1 in *C. elegans* has been shown to bind directly to PP2A-B56^38^. We therefore first investigated if the PP2A-B56 binding motif in human Bub1 was functionally important. Using our Bub1 complementation approach we observed no effects on chromosome alignment or SAC activity when we deleted amino acids 634 to 686 which encompasses the putative Bub1 PP2A-B56 binding motif (Bub1ΔB56) (Fig. 2A). Consistently, ITC measurements of Bub1 peptides encompassing the binding motif showed very weak binding to B56α (Fig. 2B). Indeed, the affinity was almost decreased 400-fold compared to the BubR1 PP2A-B56 binding motif^27^. Furthermore, immunoprecipitation experiments showed that Bub1 from mitotic cells did not interaction with PP2A-B56 (Fig. 2D and Supplemental Fig. 2A). We thus conclude that Bub1 does not encode a functionally PP2A-B56 binding motif in human cells.

**Figure 2.**
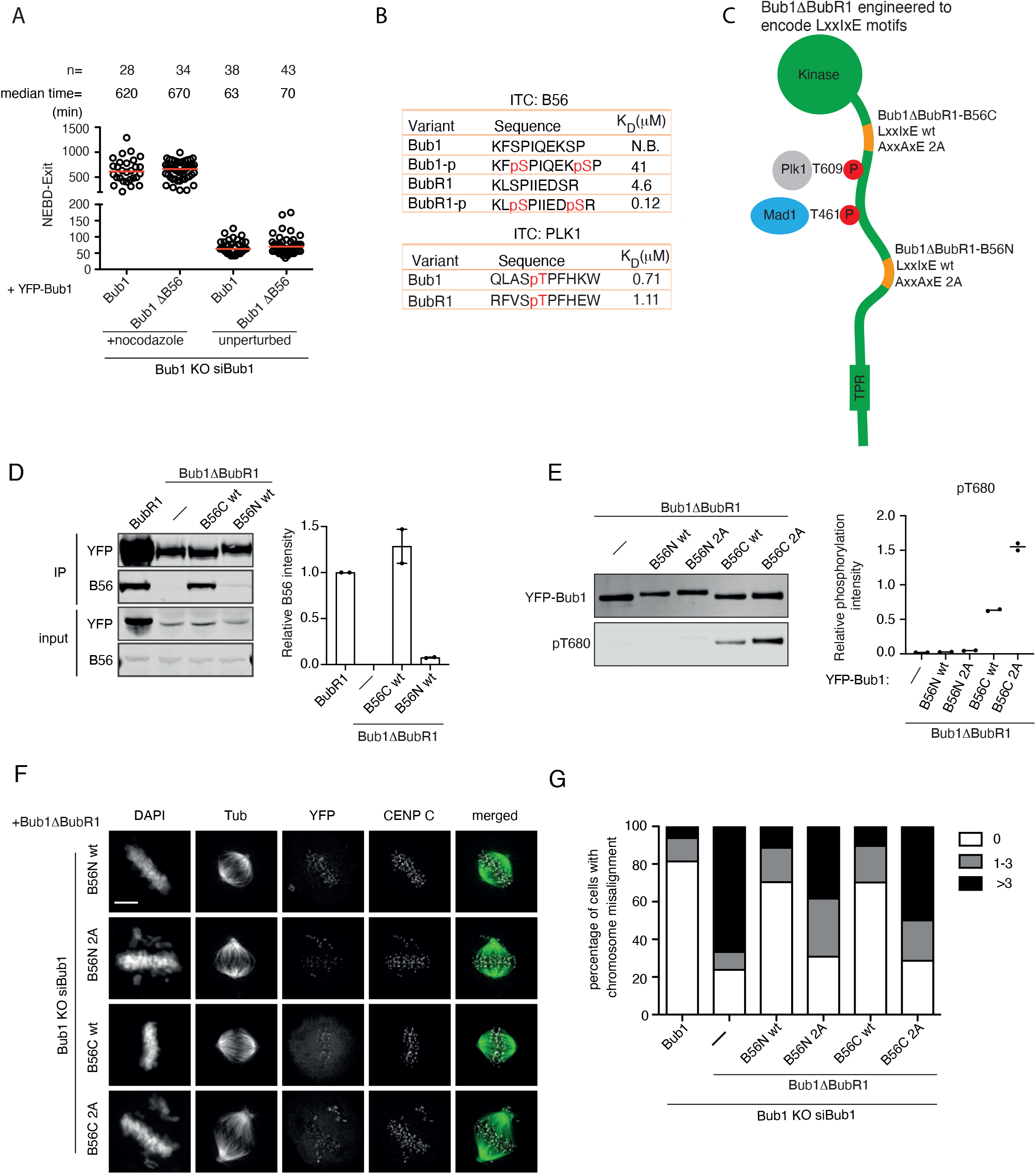
Engineered Bub1 proteins binding to PP2A-B56. **A**) The time from NEBD to mitotic exit of nocodazole treated cells or NEBD to anaphase of untreated cells complemented with the indicated YFP-tagged Bub1 constructs. Each circle represents the time a single cell spent in mitosis and the red line indicates the median time. The number of cells analysed per condition is indicated above (n=X). Representative experiment of two independent experiments. **B**) Affinity measurements of Bub1 and BubR1 peptides binding to B56α or the Plk1 polobox domain. The affinities for BubR1 peptides binding to B56α is from Kruse et al 2013. **C**) Schematic of the position of the LxxIxE motifs in the engineered Bub1 proteins. **D**) Quantitative western blot analysis of YFP immunoprecipitates of the indicated YFP-tagged proteins probed for B56α and YFP. The level of B56α was normalized to that of YFP in the precipitated samples. Quantification of two repeats is presented on the right with the level in YFP-BubR1 set to 1. **E**) Quantitative western blot analysis of the phosphorylation on BubR1 T680 in immunoprecipitates of the indicated YFP-tagged proteins. The level of T680 phosphorylation was normalized to YFP in the precipitated samples. Quantification of two repeats is presented on the right. **F-G**) Representative images of cells expressing Bub1ΔBubR1-B56N and Bub1ΔBubR1-B56C constructs after depletion of endogenous Bub1. Fixed cells were stained by DAPI and antibodies for tubulin, YFP and CENP-C. The cells were analysed by microscopy and chromosome alignment defects were determined. 100 cells were analysed per condition. Representative experiment of two independent experiments. Scale bar, 5 um.

If the main function of the Bub1-BubR1 interaction in chromosome alignment is to recruit PP2A-B56 to kinetochores one prediction would be that engineering a PP2A-B56 binding motif into Bub1ΔBubR1 would rescue chromosome alignment defects. To test this, we engrafted residues 649-697 from BubR1, which encompasses its PP2A-B56 LxxIxE binding motif into two distinct positions of Bub1ΔBubR1 (Fig. 2C). In one engineered Bub1 protein we engrafted the region into the BubR1 binding site of Bub1 (Bub1ΔBubR1-B56N). In the other, we replaced the non-functional B56 motif of Bub1 with that of BubR1 (Bub1ΔBubR1-B56C). As a control we generated the same engineered Bub1 proteins but with two amino acid mutations in the PP2A-B56 binding site (Bub1ΔBubR1-B56N/C 2A, LxxIxE mutated to AxxAxE). We first tested if these engineered Bub1 proteins bind PP2A-B56 by immunopurifying YFP-tagged Bub1 proteins from mitotic cells. We observed a strong interaction between Bub1ΔBubR1-B56C and PP2A-B56 and the amount co-purified was comparable to that co-purified with BubR1 (Fig. 2D). The Bub1ΔBubR1-B56N also bound to PP2A-B56 but to a much lesser extent. The interaction to PP2A-B56 was abolished in both Bub1ΔBubR1-B56 2A constructs (Supplemental Fig. 2A). A molecular explanation for why Bub1-B56C binds more PP2A-B56 is possibly the close proximity of the engrafted BubR1 fragment to the Plk1 docking site (T609) in Bub1^33, 39^. The interaction between PP2A-B56 and BubR1 is strongly stimulated by Plk1 phosphorylation of BubR1 S676 and T680 with Plk1 binding to T620 in BubR1^26, 27, 31^. The binding of Plk1 to T609 in Bub1 is very similar to T620 in BubR1 and since the two Plk1 binding motifs bind with similar affinity to the Plk1 polobox domain (Fig. 2B) this would likely result in efficient S676 and T680 phosphorylation in Bub1ΔBubR1-B56C. Consistent with this, we observed a high level of T680 phosphorylation specifically in Bub1ΔBubR1-B56C which was abolished by the mutation of T609 (Fig. 2E and Supplemental Fig. S2B-C).

To test if the engineered Bub1 proteins supported chromosome alignment we first analyzed this using immunofluorescence assays. In these assays we added a proteasome inhibitor hereby avoiding any indirect effects from changes to SAC signaling in the different Bub1 proteins. We scored unaligned chromosomes in individual cells which revealed that engrafting a functional PP2A-B56 binding site into Bub1ΔBubR1 supported chromosome alignment (Fig. 2 F-G). Next, we analyzed these mutants by live cell microscopy and recorded both mitotic timing and defects in chromosome alignment. In both Bub1ΔBubR1-B56N/C we observed that mitotic timing was reduced compared to Bub1ΔBubR1 and this reduction was dependent on a functional PP2A-B56 binding site (Fig. 3A-B). However, Bub1ΔBubR1-B56 N complemented cells still spent longer time in mitosis compared to Bub1 wt complemented cells. Further analysis revealed a delay in chromosome alignment in these cells whereas they only entered anaphase with fully aligned chromosomes indicating a functional SAC. In contrast, Bub1ΔBubR1-B56C complemented cells exited mitosis with a similar timing as wt complemented cells but with unaligned chromosomes suggesting failure in proper SAC signaling.

**Figure 3.**
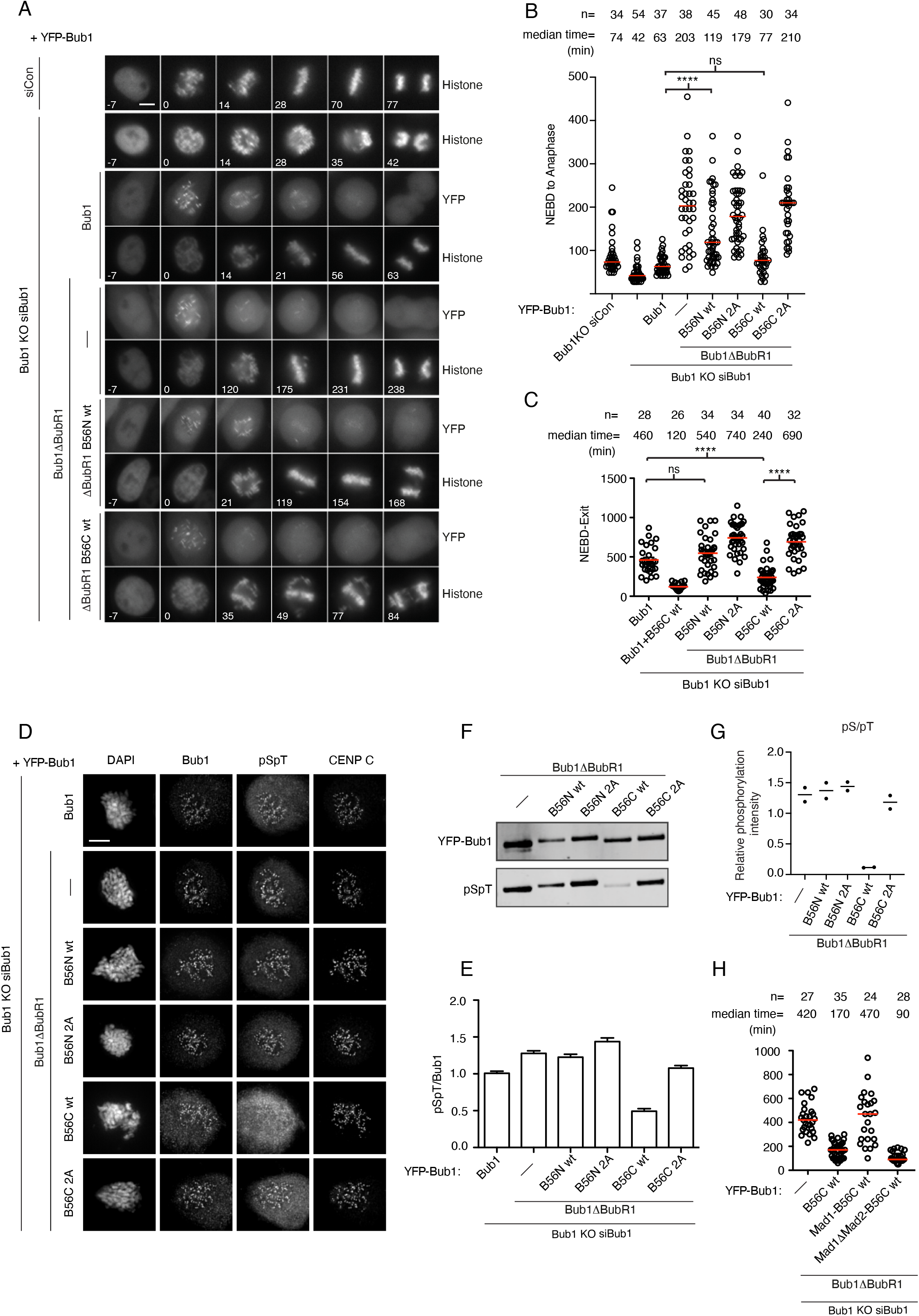
Disruption of SAC activity in Bub1 binding to PP2A-B56. **A**) Representative still images of mitosis of cells complemented with the indicated YFP-tagged constructs and expressing a CFP-tagged histone marker. Scale bar, 5 um. **B**) The time from NEBD to anaphase for the indicated conditions. Each circle represents the time a single cell spent in mitosis and the red line indicates the median time. Number of cells analysed per condition is indicated on top (n=X). Pooled data from 3 independent experiments. **C**) The time from NEBD to exit in the presence of nocodazole. Each circle represents the time spent in mitosis of a single cell. Red line indicates the median time and number of cells analysed indicated above (n=X). Representative experiment of two independent experiments. Mann-Whitney u-test was applied. ns means not significant. **** means P<0.0001. **D**) Representative images of cells complemented with the indicated YFP-Bub1 constructs and stained with DAPI and antibodies for Bub1, Bub1 pS459/pT461 (pSpT) and CENP-C. Scale bar 5 um. **E**) Quantification of the kinetochore signal of Bub1 pS459/pT461 normalized to the Bub1. At least 150 kinetochores from 10 cells were quantified and plotted for each condition. The experiment was repeated twice and the result of one experiment shown. **F-G**) Quantitative western blot analysis of Bub1 pS459/pS461 in the indicated YFP-Bub1 proteins purified from nocodazole arrested cells. The level of pSpT/YFP from two repeats is shown in G. **H**) The time from NEBD to mitotic exit in nocodazole treated cells complemented with the indicated Bub1 constructs. Each circle represents the time spent in mitosis of a single cell. Red line indicates the median time and number of cells analysed indicated above (n=X). Representative experiment of two independent experiments.

Collectively these data support the conclusion that recruitment of PP2A-B56 to Bub1 can support the alignment of chromosomes but not as efficient as when PP2A-B56 is recruited to kinetochores via BubR1.

### Recruitment of PP2A-B56 to Bub1 results in SAC failure due to dephosphorylation of the Mad1 binding site

The fact that Bub1ΔBubR1-B56C complemented cells exited mitosis with unaligned chromosomes argued for an underlying defect in SAC signaling. Indeed, challenging cells with nocodazole confirmed a specific SAC defect in Bub1ΔBubR1 B56C (Fig. 3C). This defect in SAC signaling was even more pronounced in cells complemented with a Bub1 construct (Bub1+B56C wt) that maintained its interaction with BubR1 and thus recruits PP2A-B56 both through BubR1 and the engineered LxxIxE motif in the Bub1 C position (Fig. 3C). To investigate the molecular basis for this we focused on the Bub1 phosphorylation sites critical for SAC function since we anticipated that they might be dephosphorylated by the PP2A-B56 bound to Bub1. Bub1 phosphorylation by Mps1 on T461 creates a binding site for Mad1 on Bub1 required for SAC signaling. We accessed the phosphorylation status of this site using a phospho-specific antibody (pSpT)^9^. In both immunopurifications and by immunofluorescence microscopy we observed a clear reduction in phosphorylation of this site in Bub1ΔBubR1-B56C that was dependent on PP2A-B56 binding (Fig. 3D-G). This would argue that the underlying molecular defect in SAC signaling in Bub1ΔBubR1-B56C is a disruption of the Bub1-Mad1 interaction. Indeed, if we fused Mad1 485-715 to the N-terminus of Bub1ΔBubR1-B56C (Mad1-B56C wt) we could suppress the SAC defect (Fig. 3H). The fusion of Mad1 485-715 unable to bind Mad2 did not suppress the SAC defect.

These results shows that efficient binding of PP2A-B56 to Bub1 is not compatible with a functional SAC resulting in chromosome segregation defects. However, low levels of PP2A-B56 recruitment to Bub1 as in Bub1ΔBubR1 B56N does not affect SAC signaling but these levels are insufficient for normal timing of mitotic progression.

### A chimeric BubR1 protein with Bub1 kinase activity does not support proper chromosome segregation

Our analysis of Bub1 functional domains revealed an important function of the kinase domain in chromosome alignment. In contrast BubR1 contains a pseudokinase domain that plays a role in BubR1 protein stability^40^. Given our results on engineering PP2A-B56 binding sites into Bub1 we next asked if there was a requirement for having the kinase activity restricted to Bub1. To test this, we replaced the BubR1 psudokinase domain with that of the Bub1 kinase domain in either its active or inactive form (the engineered protein encodes BubR1 1-731 fused to Bub1 734-1085 and is referred to as MadBub wt or KD as in a previous publication^23^). We investigated the ability of these fusion proteins to support mitotic functions by depleting BubR1 by RNAi and complementing cells with RNAi resistant YFP-tagged versions. Analysis of mitotic timing in unperturbed cells revealed a slight increase in MadBub wt complemented cells and this increase was dependent on a functional Bub1 kinase domain (Fig. 4A-B). This mitotic delay correlated with an increase in alignment defects arguing that MadBub wt does not efficiently support proper chromosome alignment (Fig. 4C). The effect on mitotic timing might be underestimated as we observed a slight impairment in checkpoint activity in MadBub wt cells (Supplemental Fig. S2D).

**Figure 4.**
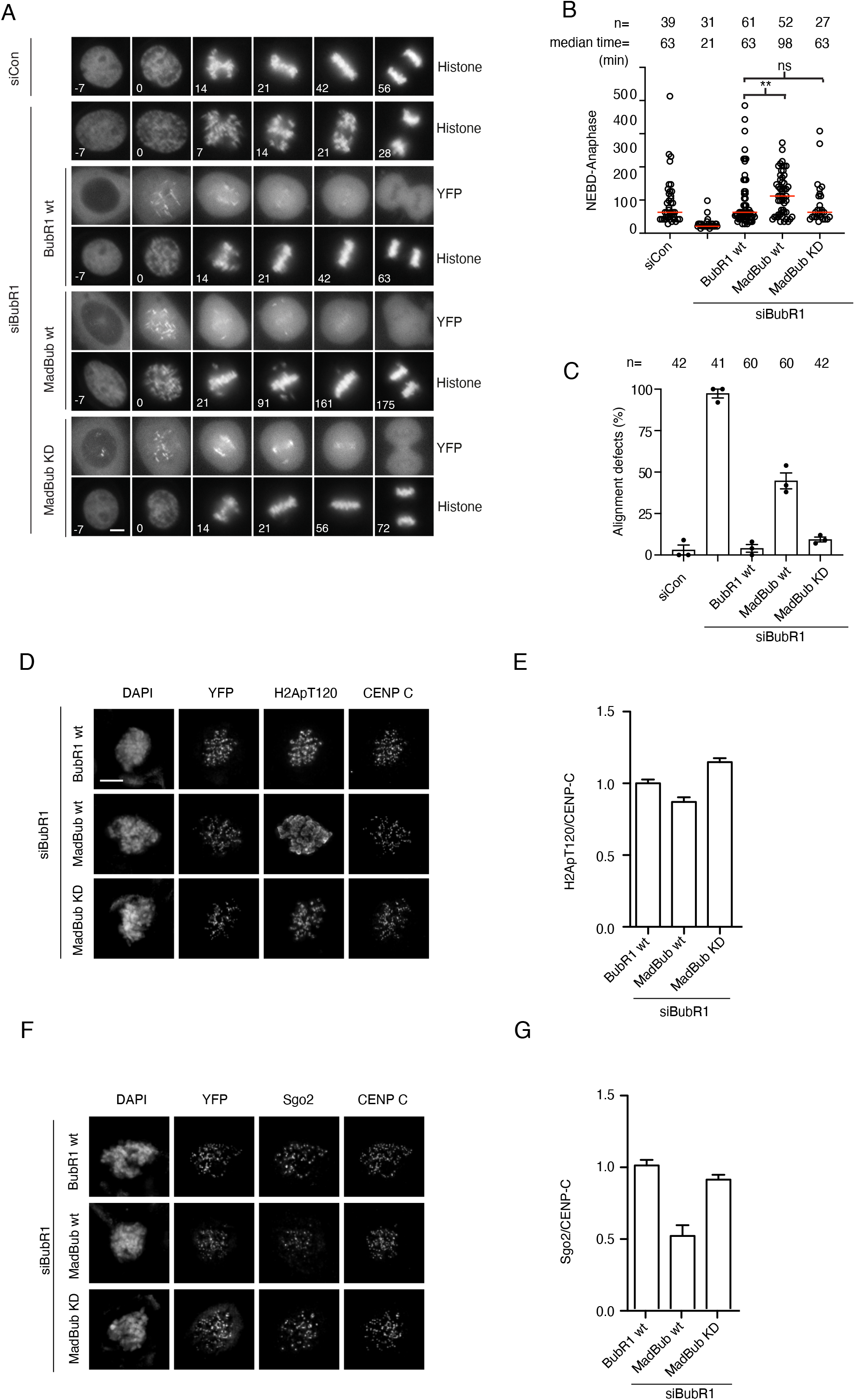
Restoring BubR1 kinase activity disrupts accurate mitosis. **A**) Representative time-lapse images of unperturbed mitosis in cells complemented with the indicated YFP-tagged BubR1 proteins and expressing a histone marker. **B**) The time from NEBD to anaphase for the indicated conditions. Each circle represents the time a single cell spent in mitosis and the red line indicates the median time. Number of cells analysed per condition is indicated on top (n=X). Pooled data from 3 independent experiments. Mann-Whitney u-test was applied. ns means not significant. ** means P<0.01. **C**) The histone marker was used to analyse alignment defects upon mitotic exit or delay in formation of metaphase plate. Each circle represents the percentage of cells with alignment defects from one experiment. Shown are results from three independent experiments. The error bars represent mean with SEM. **D**) Representative images of cells complemented with the indicated BubR1 constructs stained with DAPI and antibodies for YFP, H2ApT120, CENP-C. Scale bar, 5 um. **E**) Quantification of kinetochore/centromere levels of H2A pT120 normalized to CENP-C levels. At least 150 kinetochores from 10 cells were quantified and plotted for each condition. The experiment was repeated twice and the result of one experiment shown. **F-G**) Similar to D-E but cells stained for hSgo2.

To investigate potential molecular defects causing chromosome alignment defects in MadBub wt complemented cells we first stained mitotic cells for H2ApT120, a known Bub1 phosphorylation site. Strikingly the staining was delocalized throughout chromosomes in MadBub wt cells and no longer focused to centromeres (Fig. 4D-E). However, the total levels of centromeric H2ApT120 was not changed. As shugoshin protein localization to centromeres depends on H2ApT120 we stained cells for hSgo2 and quantified centromeric localization. We observed a 50% reduction in centromeric hSgo2 specifically in MadBub wt expressing cells (Fig. 4F-G). Although shugoshin proteins are required for proper localization of the chromosomal passenger complex (CPC) we did not observe any changes in the centromeric localization of the CPC component INCENP (Supplemental Fig 2E). Our analysis of MadBub shows that engrafting a functional Bub1 kinase domain onto BubR1 is not compatible with proper chromosome segregation which could be due to delocalized H2ApT120 impacting on proper centromeric protein localisation. Potentially our engineered MadBub as part of the diffusible MCC complex allows for delocalized phosphorylation events, which could explain why BubR1 is a pseudokinase.

Here we provide important insight into the spatial positioning of phosphatase and kinase activity within the Bub1-BubR1 complex. Our data argue that in human cells the optimal function of the Bub1-BubR1 complex in the SAC and chromosome alignment depends on precise positioning of phosphatase and kinase activity within the complex. Although recruiting PP2A-B56 directly to Bub1 at levels like BubR1 can support chromosome alignment it disrupts the phosphodependent interaction with Mad1 hereby imparing SAC activity. This likely explains why the putative B56 binding site in Bub1 is not functional. Since Bub1 T461 is dephosphorylated by PP2A-B56 bound to BubR1^41^ this argues that the LxxIxE motif in BubR1 is optimally positioned to balance the activity of the phosphatase in the SAC and chromosome alignment.

Interestingly in *C. elegans* Bub1 harbors a functional LxxIxE motif at position 282-287 C-terminal to the Bub3 binding site while in budding yeast the LxxIxE motif is present in Mad3 like human cells^28^. The fact that Bub1 in *C. elegans* contains a functional LxxIxE motif could suggest that dephosphorylation of T407, the interaction site for Mad1, is less efficiently dephosphorylated by the bound PP2A-B56^11, 38^. This would be in line with our data on Bub1 B56N that allows for a functional checkpoint as it does not affect phosphorylation of Bub1 T461. However, the levels of PP2A-B56 recruited by Bub1 B56N appears insufficient to effectively support chromosome alignment in human cells. Furthermore, additional contacts between Mad1 and Bub1 in *C. elegans* could make the interaction less dependent on T407 phosphorylation and thus allows for a functional LxxIxE motif in Bub1^8^. This would be reminiscent of the fusion of Mad1 to Bub1ΔBubR1 B56C which suppressed the SAC defect. In conclusion the precise spatial position of kinase and phosphatase activity within the Bub1-BubR1 complex is critical for balancing the chromosome alignment and SAC functions of this complex. Determining how the ancestral MadBub protein evolved to either maintain or separate its kinase and phosphatase activities in different organisms and how this balances its different functions in chromosome segregation will be interesting to explore.

## ACKNOWLEDGEMENTS

Work at the Cancer Institute is supported by grant 31970666 from the National Natural Science Foundation of China and Taishan Scholar Project tsqn201812054 from Shandong, China. Work at the Novo Nordisk Foundation Center for Protein Research is supported by grant NNF14CC0001 and JN is supported by grants from the Danish Cancer Society (R269-A15586-B17), Independent Research Fund Denmark (8021-00101B and 0134-00199B) and Novo Nordisk Foundation (NNF20OC0065098). We thank the protein production and characterization facility at NNF CPR for producing recombinant proteins. We thank Jennifer DeLuca for providing Ndc80 phosphoantibodies.

## AUTHOR CONTRIBUTIONS

LW cloned Bub1 constructs, performed live cell imaging for SAC and all the immunofluorescence analysis with the help of YZ, CS, LZ. TK performed immunopurifications and analysis of MadBub fusions and live cell imaging of Bub1. BLM performed ITC measurements. BL, JF and ZL provided scientific input and discussed the project with GZ. JN and GZ coordinated the work and wrote the manuscript together with TK.

## CONFLICT OF INTEREST

The authors have no conflict of interest

## FIGURE LEGENDS

**Supplemental figure 1.**
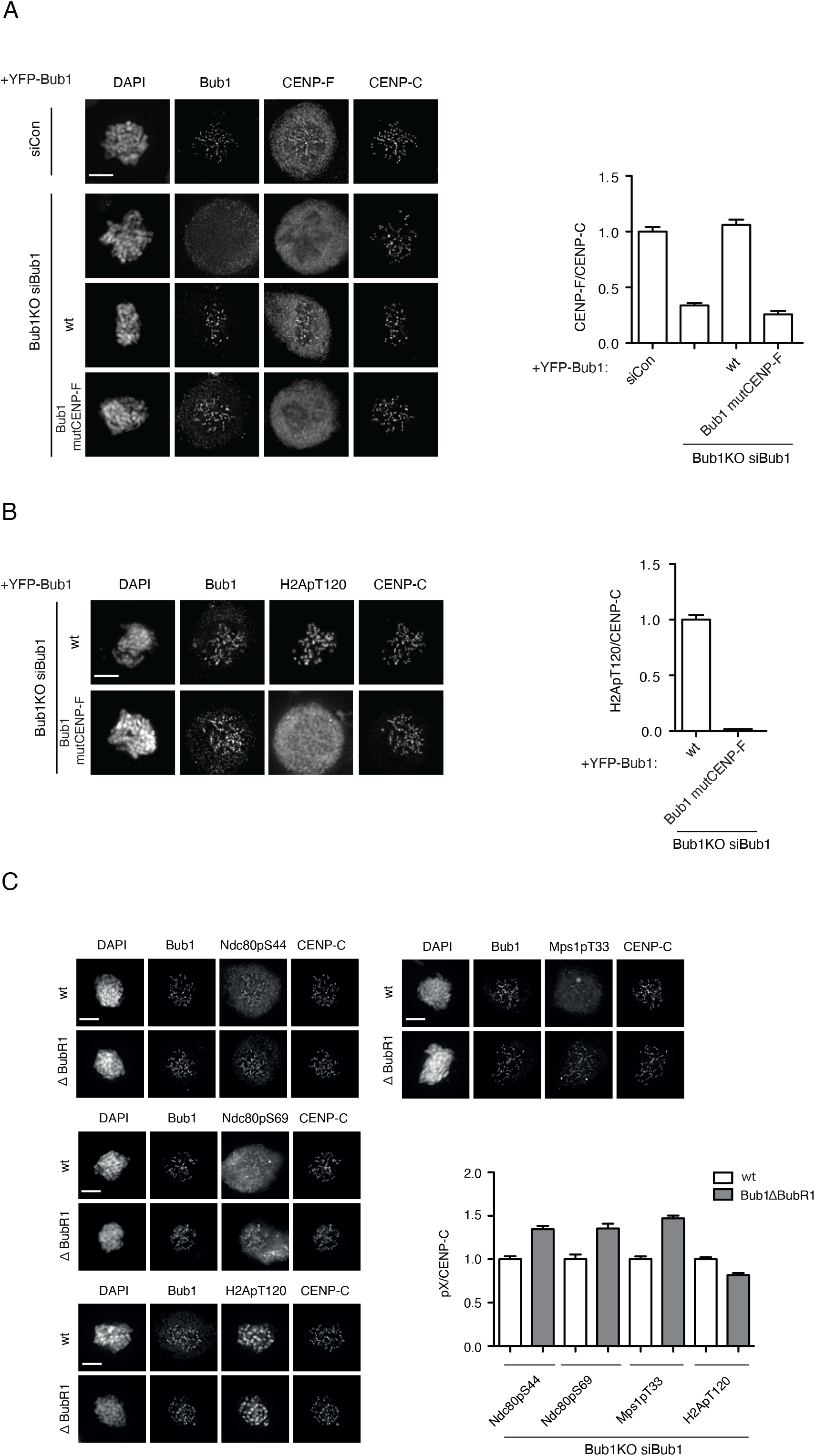
**A)** Nocodazole arrested cells complemented with the indicated YFP-tagged constructs and stained with DAPI and antibodies to Bub1, CENP-F and CENP-C. The kinetochore levels of CENP-F normalized to CENP-C is shown on the right. At least 150 kinetochores from 10 cells were quantified and plotted for each condition. The experiment was repeated twice and the result of one experiment shown. **B)** Similar to A) but cells stained for H2ApT120. **C)** Nocodazole arrested cells complemented with Bub1 wt or Bub1ΔBubR1 and stained for Bub1, CENP-C and the indicated phosphorylation specific antibodies. The kinetochore levels of the phosphospecific antibody were normalized to CENP-C. At least 150 kinetochores from 10 cells were quantified and plotted for each condition. The experiment was repeated twice and the result of one experiment shown.

**Supplemental figure 2.**
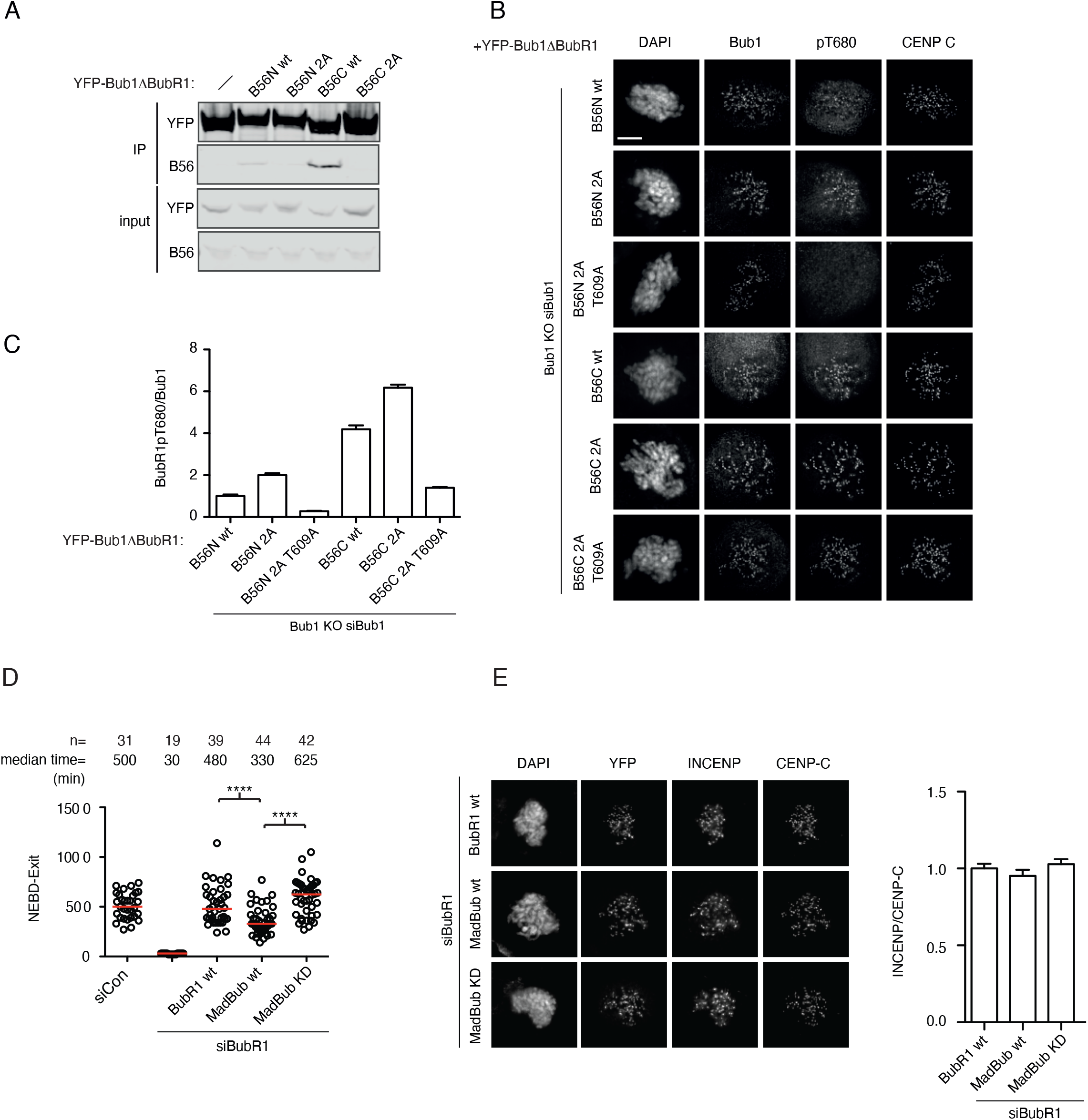
**A)** Affinity purifications of the indicated YFP-tagged proteins from nocodazole arrested HeLa cells. The purifications were stained for YFP and B56α. **B**) Representative images of nocodazole arrested cells complemented with the indicated YFP-tagged constructs and stained with DAPI and antibodies to Bub1, BuBR1 pT680 and CENP-C. **C**) The kinetochore levels of BubR1 pT680 normalized to CENP-C is shown on the right. At least 150 kinetochores from 10 cells were quantified and plotted for each condition. The experiment was repeated twice and the result of one experiment shown. **D**) Cells complemented with the indicated YFP-tagged BubR1 constructs were challenged with nocodazole and time from NEBD to mitotic exit determined by time-lapse microscopy. Each circle represents the time spent in mitosis of a single cell. Red line indicates the median time and number of cells analysed indicated above (n=X). Representative experiment of two independent experiments. Mann-Whitney u-test was applied. ns means not significant. **** means P<0.0001. **E)** Nocodazole arrested cells complemented with the indicated YFP-tagged constructs and stained with DAPI and antibodies to YFP, INCENP and CENP-C. The kinetochore/centromere levels of INCENP normalized to CENP-C is shown on the right. At least 150 kinetochores from 10 cells were quantified and plotted for each condition. The experiment was repeated twice and the result of one experiment shown.

## METHODS

### Cell culture and RNAi

HeLa or HeLa Bub1KO cells were cultivated in DMEM medium (Gibco) supplemented with 10% FBS and antibiotics. For RNAi and rescue experiments, cells were seeded in 6-well plate around 50% confluence with thymidine (2.5 mM) added in DMEM medium. 24 hours later, cells were released from thymidine arrest and transfected with siRNA oligo with RNAi-resistant constructs using Lipofectamine 2000 (Invitrogen). For Bub1 depletion, a second RNAi was performed 24 hours later in the presence of thymidine. For BubR1 depletion, thymindine was applied to the medium without a second RNAi. The following morning, cells were released from thymidine arrest and processed for further assays. RNAi oligos targeting Bub1 (5’GAGUGAUCACGAUUUCUAAdTdT3’), BubR1 5’GAUGGUGAAUUGUGGAAUAdTdT3’) or luciferase (5’ CGUACGCGGAAUACUUCGAdTdT 3’) was synthesized from Sigma and used for the RNAi.

### Cloning

The constructs used in this study were constructed using standard restriction cloning methods and mutagenesis by PCR. Briefly, wild type Bub1 or BubR1 was cloned into pcDNA5/FRT/TO N-YFP vector by KpnI and NotI. To engineer the Bub1ΔBubR1-B56N/C constructs, BamHI site was inserted separately into the Bub1ΔBubR1 cDNA at corresponding positions by PCR. The sequence encoding the region encompassing the BubR1 B56 binding motif (649-697aa) was inserted using BamHI. To generate MadBub constructs, BubR1 cDNA for N-terminal BubR1 (1-731aa) was first cloned into N-YFP vector by KpnI and BamHI followed by insertion of the sequence encoding Bub1 kinase domain (734-1085aa) by BamHI and NotI. To generate Bub1ΔCENP-F construct, a short sequence for an alternative alpha helix (5’ TCATCCGAAGAGTACGCTCGTAACTGGGCTGCACTAAAC3’) was cloned into the Bub1 cDNA to replace the sequence for the C-terminus alpha helix (1066-1082aa). The details of cloning will be provided upon request. Gene amplification or mutation PCR was performed with KOD DNA polymerase (Toyobo). All the restriction enzymes were from Thermo Scientific.

### Live cell imaging

HeLa cells were seeded in 6-well plate at confluence around 50% and synchronized with thymidine. RNA interference was performed as described in the above section. 750 ng of RNAi-resistant YFP-Bub1 or YFP-BubR1 construct and 30 ng of CFP-H3 were co-transfected in each well for undisburbed mitosis imaging. No CFP-H3 was used for SAC assay. The day before live cell imaging, cells were re-seeded into an 8-well chamber slide (Ibidi) with thymidine in the medium. On the day of live cell imaging, cells were released from thymidine 5 hours before the start of filming. Leibovitz’s L-15 medium (Gibco) containing 10% FBS was applied into each chamber before microscopy. For SAC assays, nocodazole (Sigma) was added used at 30 ng/ml. Deltavision Elite system (Cytiva) or Nikon A1HD25 imaging system (Nikon) was used for live cell imaging. YFP and CFP signals were collected every 7 or 10 minutes for a total 24 hours. Softworx or NIS-Elements AR Analysis was used for data analysis.

### Immunofluorescence, antibodies and quantification

Cells growing on coverslips were treated as described in the above. Cells were released from the second thymidine arrest into DMEM medium containing RO3306 (5 uM) for 12 hours. Afterwards, syncrhonized cells were released into fresh medium containing nocodazole (200 ng/ml) for 45 minutes or MG132 (10 uM) for 105 minutes. Cells were fixed by 4% paraformaldehyde in PHEM buffer containing 60 mM PIPES, 25 mM HEPES, pH 6.9, 10 mM EGTA and 4 mM MgSO_4_ at room temperature for 20 minutes. 0.5% Triton X-100 in PHEM was used to permeabilize the fixed cells for 10 minutes at room temperature. The antibodies used in this study include Bub1 (Abcam, ab54893, 1:200), Bub1 pSpT (home made, 1:200), CENP-C (MBL, PD030, 1:800), BubR1 pT680 (Abcam200061, 1:500), alpha tubulin (Sigma, F2168, 1:400), CENP-F (Abcam, ab5, 1:200), H2A pT120 (Active Motif, 39391, 1:400), Ndc80 pS44, Ndc80 pS69 (kind gift from Jennifer DeLuca, 1:200), Mps1 pT33 (home made, 1:200). Fluorescent secondary antibodies are Alexa Fluor Dyes (Invitrogen, 1:1000) except GFP booster-FITC was used for YFP detection (Chromotek, gb2AF488-10, 1:500). Z-stacks with 200 nm intervals were taken with Thunder Imaging System (Leica) using a 100x oil objective followed by deconvolution with Thunder cleaning function. Signal quantification was performed by drawing a circle around each kinetochore. The three continuous peak values were averaged and subtracted of the background values from a neighboring circle. At least 150 kinetochores from 10 cells were quantified and plotted in each condition. At least two repeats were performed for each experiment and the representative images were shown in figures.

### Immunoprecipitation and Western blot

HeLa cells were transfected with YFP tagged Bub1 or BubR1 constructs. 36 hours later, the cells were treated with nocodazole (200 ng/ml) for an additional 12 hours. Mitotic cells were collected by shake off and lysed in buffer containing 10 mM Tris pH 7.5, 150 mM NaCl, 0.5 mM EDTA and 0.5% NP-40. For purifications to detect PP2A-B56 binding, a low salt buffer was used as described previously^27^. After centrifugation at 16,000 g for 15 minutes, the supernatant was incubated with GFP-Trap agarose beads (ChromoTek) and shaken at 1200 rpm for 30 minutes at 4 degrees Celsius. After three washes, the bound protein was eluted in 2xLDS sample buffer and applied to SDS-PAGE for detection of indicated proteins. Antibodies used in this study include YFP antibody (home made, 1:2000), Bub1 pSpT (home made, 1:1000), BubR1 pT680 (Abcam200061, 1:1000), B56 alpha (BD, 610615, 1:1000).

### Peptide binding assay

Peptides were purchased from Peptide 2.0 Inc (Chantilly. VA, USA). The purity obtained in the synthesis was 95 – 98 % as determined by high performance liquid chromatography (HPLC) and subsequent analysis by mass spectrometry. Prior to ITC experiments both the protein and the peptides were extensively dialyzed against 50 mM sodium phosphate, 150 mM NaCl, 0.5 mM TCEP, pH 7.5 or 50 mM sodium phosphate, 150 mM NaCl, 0.5 mM TCEP, pH 7.5. All ITC experiments were performed on an Auto-iTC200 instrument (Microcal, Malvern Instruments Ltd.) at 25 °C. Both peptide and protein concentrations were determined using a spectrometer by measuring the absorbance at 280 nm and applying values for the extinction coefficients computed from the corresponding sequences by the ProtParam program (http://web.expasy.org/protparam/). The peptides at approximately 450 μM concentration were loaded into the syringe and titrated into the calorimetric cell containing the B56α at ∼ 40 μM. The reference cell was filled with distilled water. If the so-called c-value, defined as the ratio of analyte concentration in the cell to K_D_, was lower than 1, a low-c assay was performed using ∼ 25 μM B56α in the sample cell and peptide concentrations between 1.0 mM – 2.0 mM in the syringe. The peptides at approximately 450 µM or 120 µM (for submicromolar affinities) concentration were loaded into the syringe and titrated into the calorimetric cell containing the Plk1 polobox domain, respectively, at 35 µM or 10 µM

In all assays, the titration sequence consisted of a single 0.4 μl injection followed by 19 injections, 2 μl each, with 150 s spacing between injections to ensure that the thermal power returns to the baseline before the next injection. The stirring speed was 750 rpm. Control experiments with the peptides injected in the sample cell filled with buffer were carried out under the same experimental conditions. These control experiments showed heats of dilution negligible in all cases. The heats per injection normalized per mole of injectant *versus* the molar ratio [peptide]/[protein] were fitted to a single-site model. A 1:1 stoichiometry complex was assumed for the fitting of the ITC binding isotherms in the case of the low c-assays. Data were analysed with MicroCal PEAQ-ITC (version 1.1.0.1262) analysis software (Malvern Instruments Ltd.).

### Protein purification

B56α was expressed and purified as previously described^27^. The Plk1 polo box domain (residues 367-603) was expressed in BL21(DE3) cells overnight at 18 degrees. Cells were harvested by centrifugation and resuspended in buffer L (50 mM NaP pH=7.5, 300 mM NaCl, 10 mM imidazole, 10% glycerol, 0.5 mM TCEP, 1XComplete EDTA free tables (Roche). Lysis was done using a high pressure homogenizer (Avestin) and lysate clarified by centrifugation. The clarified lysate was applied to a 5ml HiTrap Nickle column and proteins eluted with a 10-500 mM imidazole gradient and peak fractions collected. Subsequently TEV protease was added to remove tag and protein dialysed into buffer L lacking imidazole and loaded onto a 5ml HiTrap Nickle column and unbound protein collected and pooled. The untagged Plk1 polo box domain was finally loaded on a superdex 200 16/60 column equilibrated with buffer G (50 mM NaP, 150 mM NaCl, 10% glycerol, 0,5 mM TCEP) and peak fractions collected.

